# LptB_2_FG is an ABC transporter with Adenylate Kinase activity regulated by LptC/A recruitment

**DOI:** 10.1101/2021.07.08.451440

**Authors:** Tiago Baeta, Karine Giandoreggio-Barranco, Isabel Ayala, Elisabete C. C. M. Moura, Paola Sperandeo, Alessandra Polissi, Jean-Pierre Simorre, Cedric Laguri

## Abstract

<u>L</u>ipo<u>p</u>oly<u>s</u>accharide (LPS) is an essential glycolipid covering the surface of gram-negative bacteria. Its transport involves a dedicated 7 protein transporter system, the Lpt machinery, that physically spans the entire cell envelope. LptB_2_FG complex is an ABC transporter that hydrolyses <u>A</u>denosine <u>T</u>ri<u>p</u>hosphate (ATP) to extract LPS from the inner membrane (IM). LptB_2_FG was extracted directly from IM with its original lipid environment by Styrene-Maleic acids polymers(SMA). SMA-LptB_2_FG in nanodiscs displays ATPase activity and a previously uncharacterized <u>A</u>denylate <u>K</u>inase (AK) activity. It catalyzes phosphotransfer between two ADP molecules to generate ATP and AMP. ATPase and AK activities of LptB_2_FG are both stimulated by the interaction on the periplasmic side with LptC and LptA partners and inhibited by the presence of LptC transmembrane helix. Isolated ATPase module (LptB) has weak AK activity in absence of LptF and LptG, and one mutation, that weakens affinity for ADP, has AK activity similar to that of fully assembled complex. LptB_2_FG is thus capable of producing ATP from ADP depending on the assembly of the Lpt bridge and the AK activity might be important to ensure efficient LPS transport in fully assembled Lpt system.

## Introduction

Gram-negative bacteria possess a double membrane system delimiting an aqueous space, the periplasm. While the cytoplasmic inner membrane is a canonical phospholipid bilayer, the outer membrane (OM) is asymmetric with its outer layer mostly composed of the <u>L</u>ipo<u>p</u>oly<u>s</u>accharide (LPS) glycolipid (1). These molecules are surface exposed, contribute to the high impermeability of the OM, and play a role in immune response, pathogenesis, and drug resistance (2). Maintenance of the OM asymmetry is essential for bacterial viability and a continuous flow of LPS molecules needs to cross the periplasm and reach the external layer to keep up with cell growth. A system of 7 essential proteins, the <u>L</u>ipo<u>p</u>olysaccharide <u>T</u>ransport System (Lpt), is dedicated to trafficking LPS across the cell envelope. These proteins (LptA-G) are found in all cellular compartments (IM, periplasm and OM) and associate through their jellyroll domains to form a continuous periplasmic bridge connecting IM and OM(3, 4). LptB_2_FG inner membrane complex is an ABC transporter in which LptB_2_ represents the Nucleotide Binding Domain (NBD) component which binds and hydrolyses <u>A</u>denosine <u>T</u>ri<u>p</u>hosphate (ATP) to generate mechanical force to transport LPS. Upon ATP binding, LptB dimerizes in a closed structure, and ATP hydrolysis relaxes the dimer into an open conformation. Conformational changes are transmitted to LptFG through coupling helices, and allow LPS extraction (5). LptB_2_FG associates with LptC via 2 distinct interactions: i) LptF/LptG interact with LptC transmembrane domain, regulate LPS entry into the complex and LptB_2_FG ATPase activity (5), ii) LptF and LptC jellyroll domains associate in the periplasm. LptA jellyroll then bridges to LptD/E OM complex that ultimately assembles the LPS to the OM outer leaflet (3).

The superfamily of <u>A</u>TP-<u>b</u>inding <u>C</u>assette (ABC) transporter proteins comprises more than 500 members, and support traffic of metabolites and molecules through ATP hydrolysis (6). Several prokaryotic ABC transporters (MsbA, TmrAB and LmrA) couple another reaction in their ATPase domain, Adenylate Kinase (AK) (7). AK catalyzes a phosphotransfer reaction between 2 <u>A</u>denosine <u>D</u>i<u>p</u>hosphate (ADP) molecules producing ATP and AMP (Adenosine mono-phosphate) with no energy consumption (8). MsbA in particular translocates LPS across the inner membrane prior to its transport by LptB_2_FG (9, 10). The additional active site for the AK has been suggested to be located proximal to the ATP binding site, since the reaction requires both ADP molecules close by in space (11).

LptB_2_FG was extracted directly from the *Escherichia coli* inner membrane with Styrene-Maleic Acids (SMA) polymers without the use of detergents. ^1^H NMR on nucleotides showed the SMA-LptB_2_FG complex has ATPase activity but also Adenylate Kinase (AK) activity, and both activities are regulated by the assembly of LptB_2_FG with LptC and LptA partners. Point-mutations were introduced in LptB_2_FG complex and in isolated LptB to address their effect on ATPase and AK activity by NMR and locate the AK site.

## Results

### LptB_2_FG extracted from the inner membrane by SMA polymers displays ATPase and Adenylate Kinase activity

LptB_2_FG complexes from several organisms have been expressed and purified in either detergent micelles (5, 12–15), reconstituted nanodiscs (5) or liposomes (16). These protocols all involve solubilization and purification of LptB_2_FG complex in detergent micelles. The use of detergent free protocols in preparation of membrane protein, to maintain their original lipid environments, becomes increasingly important to ensure functionality and stability. In that aspect the use of SMA polymers and its derivatives, that allow direct extraction of membrane proteins from purified membranes, provides a versatile tool to study membrane proteins as close as possible to their natural environment (17, 18). LptB_2_FG complex was extracted directly from the inner membrane of *Escherichia coli* using SMA polymers. SMA-LptB_2_FG was successfully purified in nanodiscs of about 10 nm diameter with the expected 2:1:1 stoichiometry (Fig. S1). LptB_2_FG ATP hydrolysis was checked in presence of LptC and LptA, its periplasmic partners, using NMR to probe the reaction (Fig. 1A). The monomeric version of LptA (LptA_m_) and the soluble version of LptC (ΔTM-LptC), which are able to sustain activity and cell viability (19, 20), were added to SMA-LptB_2_FG and ATP hydrolysis was followed by real-time ^1^H-NMR (Fig. 1B). NMR is a spectroscopic technique that provides resonance frequency of active nuclei, in our case protons, depending on their chemical environments (21). It can discriminate between ATP and ADP as several ^1^H shift frequencies upon phosphate loss, here hydrogen H4’ on the ribose (Fig. 1B). SMA-LptB_2_FG shows ATP hydrolysis, with fast disappearance of peaks characteristic of ATP and concomitant appearance of ADP specific peaks in the NMR spectrum. The rate of ATP hydrolysis is 6.5 moles of ATP consumed/min/mole of LptB. After the initial rapid accumulation of ADP due to ATPase activity, ADP level decreases at the same rate as the appearance of a new set of NMR peaks (8.60 ppm in the H8 region, and 4.39 ppm in H4’ region). This suggests conversion of ADP into another specie over time and its ^1^H NMR chemical shifts suggest it could correspond to AMP (22). ^31^P NMR spectrum was collected at the end of the reaction and showed the characteristic peak of AMP at 3 ppm (Fig. S2). Conversion of ADP into AMP is compatible with an Adenylate Kinase reaction 2ADP ⇔ ATP + AMP, already been observed for several ABC transporters (7, 11). ATP was replaced by ADP as a substrate and reaction was followed in identical conditions in real time (Fig. 1B). ADP decrease is seen immediately, together with appearance of AMP and ATP. Generation of ATP is consistent with AK activity and excludes hydrolysis of ADP into AMP. ATP levels observed are lower than those of AMP, as AK and ATPase activities occur simultaneously, and newly generated ATP is partly hydrolyzed back to ADP. The initial rates of AK activity are much slower than ATPase with 0.2 and 0.7 moles of AMP produced/min/mole of LptB with ATP or ADP as starting substrates (Fig. 1B).

**Figure 1-.**
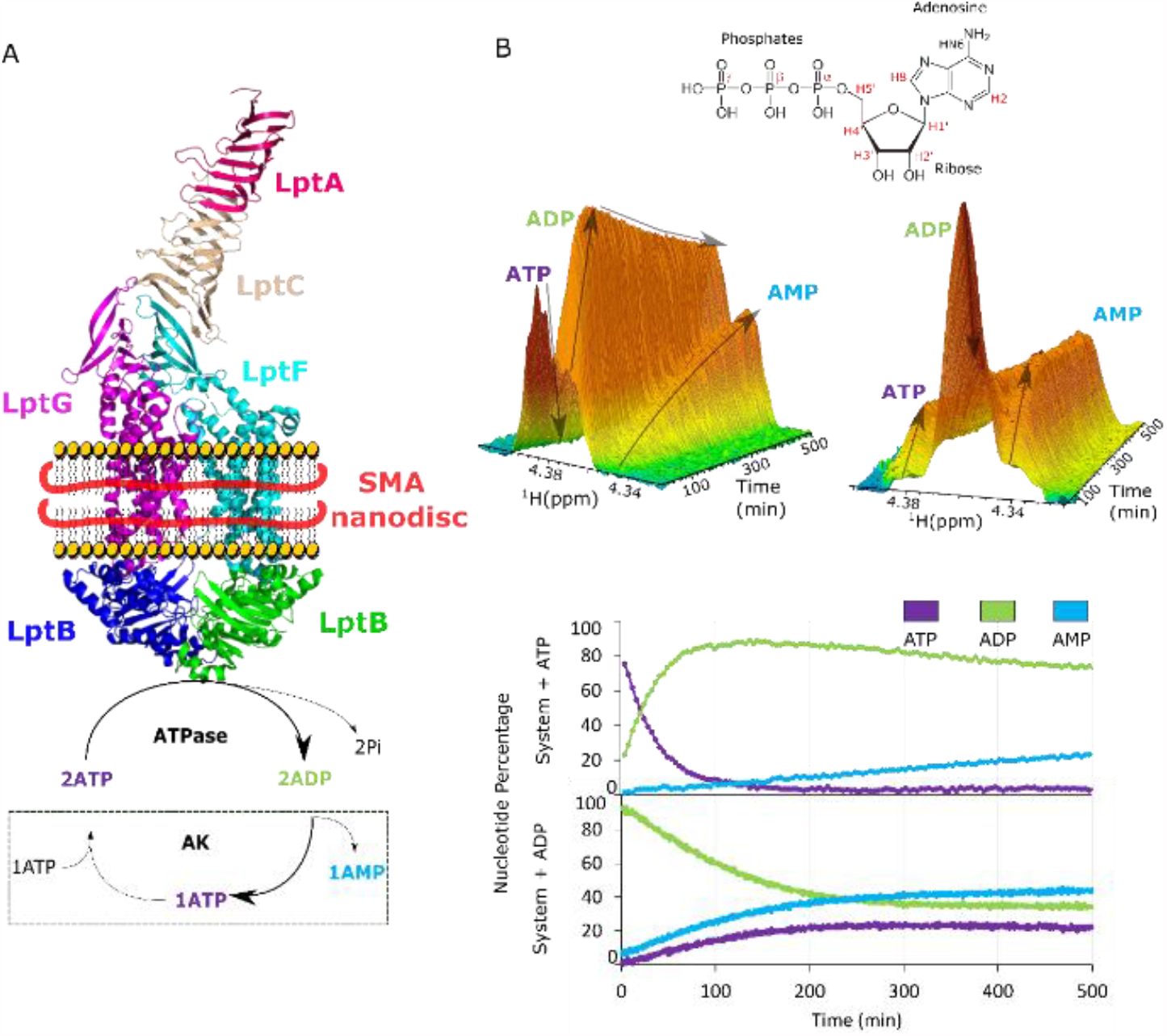
(A) Representation of LptB_2_FGCA (with ΔTM-LptC and LptAm) system in SMA nanodiscs with its updated enzymatic cycle. (B) Real-time kinetics of LptB_2_FGCA observed by ^1^H-NMR with either ATP (left) or ADP (right) as substrate over time with evolution of the characteristic H4’ peaks of ATP, ADP and AMP. Nucleotide quantification along the experiment is represented (bottom).

### ATPase and AK activities are regulated by the assembly of the Lpt system

LptB_2_FG ATPase activity is repressed by the TM domain of LptC (5, 16), we thus examined ATPase and AK activities of SMA-LptB_2_FG with/without ΔTM-LptC/LptA_m_ and in complex with full length LptC with/without LptA_m_ (Fig. 2A and 2B). For the following experiments nucleotides were quantified by NMR at the end of incubation (Fig. S3). SMA-LptB_2_FG displays less ATPase and AK activity in absence of ΔTM-LptC and LptA_m_. Proper assembly of the periplasmic part of LptC and its complex with LptA (20) is thus important in stimulating both ATPase and AK activity. SMA-LptB_2_FGC, containing full length LptC has no ATPase activity, but the repression is relieved in presence of LptA_m_ (Fig. 2B). Similarly, SMA-LptB_2_FGC has little AK activity (Fig. 2B) which is increased by addition of LptA_m_. AK activity and ATPase activities are both stimulated by the assembly of LptB_2_FGCA complex without the transmembrane LptC segment. In intact LptB_2_FGC ATPase and AK are activated by assembly with LptA.

**Figure 2-.**
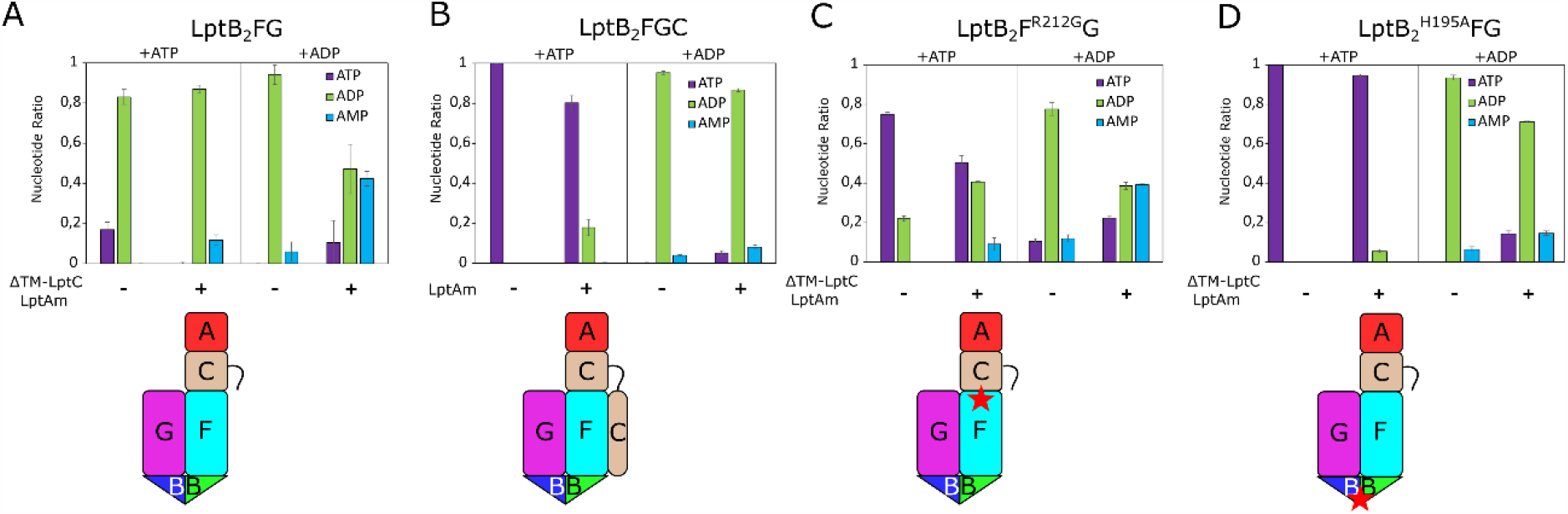
(Top Panel) **Assembly on the periplasmic or membrane level with LptC and LptA influence ATPase and AK activity of LptB**_**2**_**FG**. SMA-LptB_2_FG is incubated with ATP or ADP and nucleotides levels (ATP, ADP and AMP) quantified by ^1^H NMR from two or three independent experiments. (A) SMA-LptB_2_FG without/with ΔTM-LptC/LptA_m_, (B) SMA-LptB_2_FGC complex without/with LptA_m_. (C and D) SMA-LptB_2_F^R212G^G and SMA-LptB_2_^H195A^FG without/with ΔTM-LptC/LptA_m_. A schematic representation of the complexes used is shown with stars marking the location of mutations.

### Variants in periplasmic LptF domain and cytoplasmic LptB differently affect AK activity

Number of mutations in LptB_2_FG were studied for their effect on the complex activity, either biochemically or for their *in vivo* functionality (23, 24). Two SMA-LptB_2_FG complexes carrying already characterized mutations were purified, one on LptF periplasmic side, and one in the ATPase domain(LptB). LptF^R212G^ variant can overcome the absence of the essential LptC protein and allow cell viability (25). R212 is located in LptF periplasmic domain at the interface with LptC and is in the path of the LPS flow(15). SMA-LptB_2_F^R212G^G has impaired ATPase activity when compared to wild-type (wt) complexes (Fig. 2C), and ΔTM-LptC and LptA_m_ stimulate ATPase and AK activity (Fig. S3). With ADP as substrate SMA-LptB_2_F^R212G^G with and without ΔTM-LptC and LptA_m_, show similar production of AMP as wt complexes (Fig. 2C). R212G mutation, while impairing the ATPase, does not affect the AK, suggesting different regulation mechanisms of the two activities.

LptB H195 is involved in γ phosphate binding of ATP (Fig. 3A) and LptB^H195A^ variant has decreased ATPase activity in isolated LptB and is deleterious for cell growth (23). SMA-LptB_2_^H195A^FG has almost knocked out ATPase activity, even in presence of ΔTM-LptC and LptA_m._ AK activity is still activated by ΔTM-LptC/A_m_ but significantly decreased compared to wild type complex (Fig. 2D, Fig. S3). Alterations in the ATPase binding site of LptB affect ATPase activity and AK activity, but the latter to a lesser degree.

**Figure 3-.**
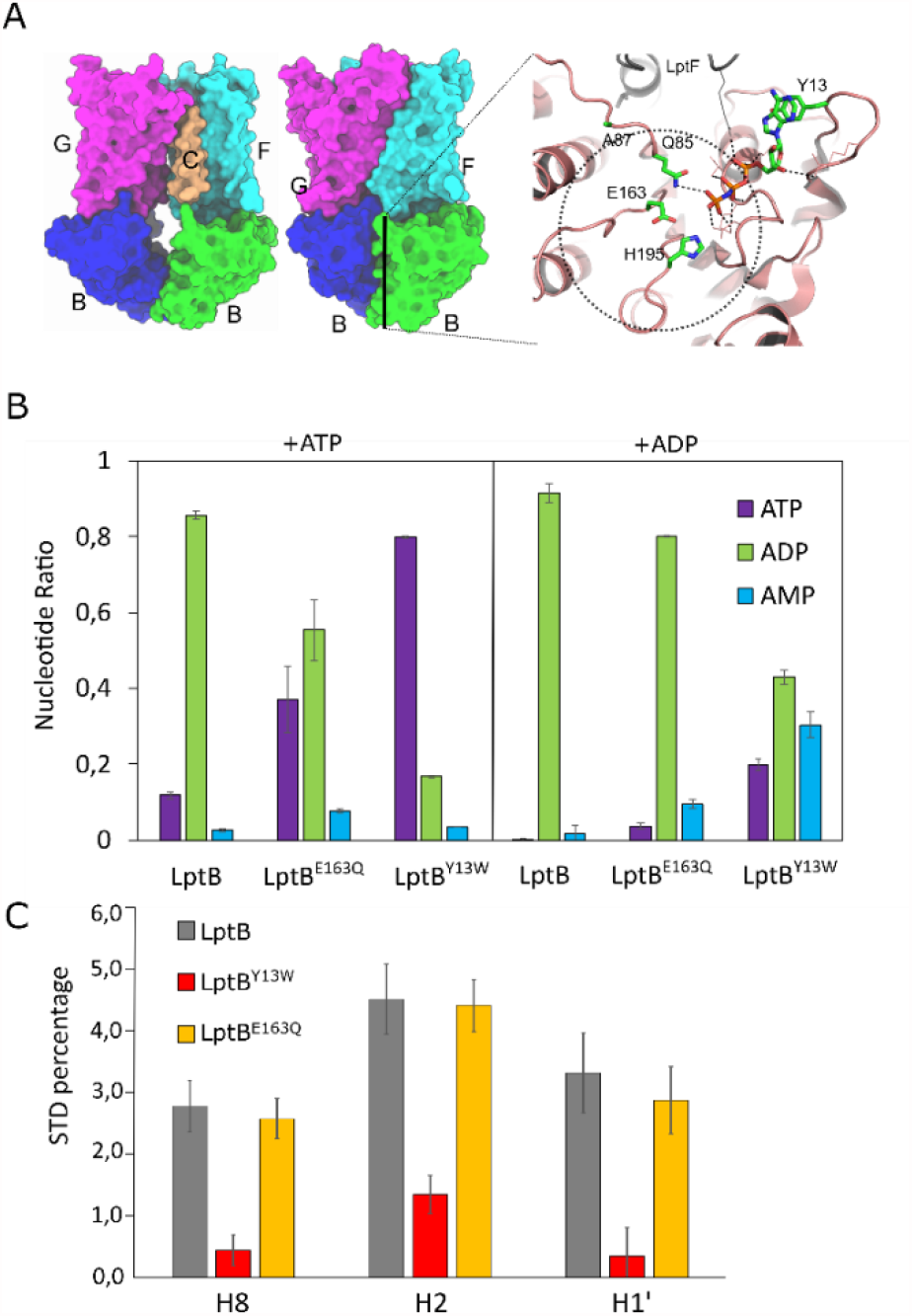
A) Structures of LptB_2_FGC in open (apo form PDB 6S8N) and closed conformation (with bound ATP analog PDB 6S8G). The jellyroll domains were not resolved in these structures as well as the LptC transmembrane helix in the open conformation. Right panel shows ATPase site with putative location of the second ADP in the AK reaction (dashed line area). Residues mutated in this study are shown (B) ATPase/AK of LptB_2_ and LptB_2_ ^E163Q^ and LptB_2_ ^Y13W^ mutants, with ATP or ADP as substrate. Nucleotide levels (ATP, ADP and AMP, in colour-code) were detected by^1^H-NMR experiment in 2 independent experiments; (C) <u>S</u>aturation <u>T</u>ransfer <u>D</u>ifferent (STD) experiments of LptB_2_ and LptB_2_^E163Q^ and LptB_2_ ^Y13W^ on ADP-βS. The three resonances shown are from H8, H2 and H1’ on the adenosine of ADP-βS.

### Mutations in the A- and B-loops of LptB stimulate AK

The location of the AK site is shared with the ATPase site (26–28). One ADP molecule occupies the ATP binding site while the other must be presented «in line» with the first one for the phosphotransfer to occur. Current models for AK suggest the second ADP molecule is located around the Q-loop (11) (Fig. 3A). The location of the second ADP molecule was probed in LptB using mutagenesis. In addition to H195A mutation in the H motif (or switch region), other mutations were inserted in canonical ABC motifs, specifically at the A-loop (Y13W), B-loop (E163Q and 163A), and Q-loop (A87Q) (Fig. 3A). Purified soluble LptB wild-type and variants proteins were incubated with either ATP or ADP, and nucleotides were quantified by ^1^H-NMR at endpoint of experiment (Fig. S4). LptB, similarly to SMA-LptB_2_FG complex (without ΔTM-LptC and LptA_m_) displays ATPase activity and little AK activity (Fig. S5). All mutations introduced in LptB decrease ATPase activity and E163A is catalytically fully inactive with no ATPase or AK activity. Two LptB variants show increased AK activity, E163Q and more significantly Y13W, the latter accumulating AMP at levels similar to SMA-LptB_2_FG/ΔTM-LptC/LptA_m_ (Fig. 3B).

To assess how binding of nucleotides relates to the activity of LptB, <u>S</u>aturation <u>T</u>ransfer <u>D</u>ifference experiments were recorded on wild-type (wt) and LptB variants. STD is an NMR technique that allows observation of the free ligand in transient binding to a protein (29). The protein is selectively saturated, and part of the saturation is transferred to the ligand during binding and a decrease of intensity is observed on the free ligand. STD on ADP by wild-type LptB shows saturation on protons H2, H8 and H1’ of the ADP, showing significant interaction with LptB. LptB^E163Q^ variant shows no difference of STD with respect to wt. STD measurement of Y13W variant with ADP is not possible, since AK activity is too fast relative to the experimental time. ADP-βS, which is not a substrate of LptB, showed STD similar as ADP with wild-type and LptB^E163Q^ variant (Fig. S6). Y13W, on the other hand, showed a highly reduced STD (Fig. 3C Fig. S6-S7). A reduction in STD can be ascribed to a reduced affinity of Y13W for ADP-βS, with a reduced residence time of the ligand, or on the contrary to a reduction in k_off_ leading to less saturation of the free ligand. A Y-W mutation in this position has already been found to reduce ATPase activity and ATP binding in other ABC transporters (30, 31). We thus conclude that LptB^Y13W^ variant increase in AK activity can be assigned to a reduction in the affinity for ADP, likely in the canonical ATPase binding site considering the position of Y13(Fig. 3A).

## Discussion

Understanding bacterial mechanisms that coordinate envelope assembly is critical to harness ways to challenge persistent and clinically relevant infections. Here, LptB_2_FG and LptB_2_FGC complexes were extracted directly from *E. coli* inner membrane in their natural lipid context, without the need for detergent extraction. LptB_2_FG complex ATPase activity is stimulated by interaction on the periplasmic side of LptB_2_FG with LptC and LptA and fully repressed when full-length LptC is coextracted with LptB_2_FG. SMA-LptB_2_FG LptB subunit is also an Adenylate Kinase. This activity is marginal until LptB_2_FG interacts with the jellyroll domains of LptC and LptA, thus forming an almost complete Lpt bridge (missing LptD/E OM complex). Establishment of the bridge must be transmitted by conformational changes through the lipid bilayer to the cytoplasmic LptB and, activate both ATPase and AK activity. LptB_2_FGC complex that contains full-length LptC with its N-terminal transmembrane domain has no ATPase activity and little AK activity until it interacts with LptA which increases both activities. Level of activity of LptB_2_FGC is much lower due to the presence of the transmembrane LptC domain that is believed to regulate LPS entry into the complex and possibly synchronizes LPS entry and ATP hydrolysis (5). The Lpt system possesses several mechanisms that regulate its activity to ensure efficient LPS transport when the Lpt system is assembled. A mutant discovered in LptF periplasmic jellyroll domain which can complement ΔLptC cells is particularly interesting. SMA-LptB_2_F^R212G^G ATPase activity is highly reduced compared to wild-type complex while AK activity is identical. This suggests that the regulation of the ATPase and AK activities by the assembly of the Lpt system are somewhat different. Nevertheless, the regulation mechanisms are currently not characterized at the molecular level.

Adenylate kinase reaction involves 2 ADP molecules, one located in the ATPase canonical site and one located nearby. While our mutagenesis study could not pinpoint the location of the binding site for the second ADP, a model of the possible locations of that ADP was determined. The complex between LptB and ADP (PDB 4P32) was used as a template, while the second ADP was docked using HADDOCK (32), with a single distant constraint between the two β-phosphates of reacting ADPs (see Methods section) (Fig. S8 Table S1). The best two clusters of models show the second ADP molecule located around Q85, E163 and H195. Q85E mutant could not be expressed, while LptB^E163Q^ showed increased AK activity, and H195A in the full complex showed reduced AK activity. While those residues could be important for the AK activity, they are all involved in the ATPase reaction, either by binding ATP γ-phosphate, the magnesium ion or coordinating water molecules (23). The models do not show extensive interactions of the second ADP with LptB suggesting, on the contrary to pure Adenylate Kinase (26), that there is no well-defined binding site for the AK activity in LptB, apart from the canonical ATP binding site. The high AK activity of LptB^Y13W^ variant cannot be easily explained. The consequence of this mutation is to strongly reduce STD, likely due to decreased affinity/residence time of the ligand in the canonical ATPase site. This could in turn accelerate the turnover of the reaction but we cannot exclude that this variant in isolated LptB somehow mimics the conformation of LptB_2_FG in presence of the jellyroll domains of LptC and LptA.

The role of the AK activity in LptB_2_FG ABC transporter (and the related MsbA) remains unknown. It is clear from results from Sherman et al. with H195A and E163Q mutations, which cannot complement Δ*lptB E. coli* cells (23), that AK activity alone without ATPase cannot sustain *E. coli* growth. Nevertheless, AK activity can generate ATP which can in turn be hydrolysed and is likely to allow LPS transport. The rate of AK activity measured in our conditions is more than ten times lower than ATPase, and the relationships between AK, ATPase and LPS transport will have to be further examined to understand the exact role of AK in LptB_2_FG.

### Experimental Procedures

#### Used Plasmids

All plasmids used were from previous studies, pET22-43 LptB(23), pCDF-Duet1-LptB_2_FG(33), pBAD/HisA-LptC(33), pQESH LptC_Δ[1-23]_ (ΔTM-LptC)(19) and LptA_Δ160_ (LptA_m_)(34). Point mutations were performed by Genewiz.

#### Strains used and Protein Expression

LptB expression was done in *E. coli* BL21 (DE3) strain (Novagen), and LptB_2_FG in *E. coli* C43 (DE3) strain (Novagen). Co-expression of LptB_2_FG and LptC was done in *E. coli* KRX cell as described (33). LptA_m_ and ΔTM-LptC were expressed and purified in *E. coli* BL21 (DE3) as described (20). For LptB and LptB_2_FG, bacterial cells were grown in Luria Broth (LB), supplemented with the correct antibiotic (ampicillin 100 µg ml^-1^ and spectinomycin 50 µg ml^-1^) at 37°C, until optical density of 600 nm (OD600) around 0.7. For both sets of proteins, induction was done with isopropyl β-D-thiogalactopyranoside (IPTG): for LptB_2_ proteins with 0.1 mM at 20°C for 16 h, while for LptB_2_FG proteins with 0.5 mM at 37°C for 3 h. Cells were harvested by centrifugation at 6000 xg for 20 min at 4°C, and frozen at -20°C until purification. Co-expression of LptB_2_FG with LptC and cell harvesting was done as before, except induction which was done with 0.02% (w/v) L-Rhamnose and 0.02% (w/v) L-Arabinose.

#### Purification of LptB_2_ proteins

Cells were mixed with buffer A containing 20 mM Tris-HCl, 150 mM NaCl, 20% (v/v) glycerol, pH 8.0, 0.5 mM Tris(2-CarboxyEthyl)Phosphine [TCEP]) and cOmplete™ EDTA-free (Sigma) and lysed by sonication. Soluble fraction was separated by centrifugation at 10.000 xg for 20 min at 4°C and loaded after addition of 10mM imidazole into Ni-NTA Agarose (QIAGEN). Resin was washed with buffer A with 20 mM Imidazole and eluted with buffer A with 300 mM Imidazole. LptB was then purified on a HiLoad^®^ 16/600 Superdex^®^ 200 pg column (GE Healthcare) in Tris-Buffered Saline (TBS) (50 mM Tris-HCl, 150 mM NaCl, pH 8.0) supplemented with 0.5 mM TCEP. LptB was concentrated with a 10 kDa cut-off Amicon^®^ Ultra Centrifugal Filter (Merck) and sample concentration determined by running a 15% SDS-PAGE with known concentration samples of <u>B</u>ovine <u>S</u>erum <u>A</u>lbumin (BSA).

#### Purification of LptB_2_FG and LptB_2_FGC

Cells in lysis buffer (50 mM Tris-HCl, 300 mM NaCl, 1 mM MgCl_2_, pH 8.0 and cOmplete™ EDTA-free (Sigma)), were lysed on a microfluidizer at 15000 psi. Cell debris are removed by centrifugation at 10000 xg for 20 min at 4°C, and membranes collected by centrifugation at 100.000 xg for 1 h at 4°C. Membranes are resuspended in 50 mM Tris-HCl, 250 mM NaCl, pH 8.0 with 0.5% SMALP 25010P (Orbiscope) for 17 h at Room Temperature (RT). Soluble SMALP particles were obtained by ultracentrifugation at 100.000 xg for 30 min at 4°C and loaded into a HisTrap™ 1 ml column equilibrated in 20 mM Tris-HCl, 150 mM NaCl, 30 mM Imidazole, pH 8.0. Elution was done in a gradient with the previous buffer supplemented with 170 mM Imidazole. Fractions containing the proteins were dialyzed against TBS buffer (20 mM Tris, 150 mM NaCl, pH 8.0) at RT and concentrated with a 100 kDa cut-off Amicon^®^ Ultra Centrifugal Filter (Merck). Sample concentration was determined by running a 15% SDS-PAGE with known concentrations of BSA.

#### Electron Microscopy

SMA-LptB_2_FG at 59µg/ml was prepared by Negative Stain <u>M</u>ica-carbon <u>F</u>lotation <u>T</u>echnique (MFT). Samples were absorbed to the clean side of a carbon film on mica, stained with 2% Na_4_O_40_SiW_12_ in distilled water, then transferred to a 400-mesh copper grid. Images were taken under low dose conditions (<10 e^-^/Å^2^) with defocus values between 1.2 and 2.5 μm on a Tecnai 12 LaB6 electron microscope at 120 kV accelerating voltage using CCD Camera Gatan Orius 1000.

#### NMR spectroscopy

Experiments were recorded on Bruker 600, 700, 850 and 950 MHz spectrometers equipped with triple ^1^H, ^13^C, ^15^N resonance cryoprobes, at 25°C for ^1^H and ^31^P, and at 37°C for real-time kinetic analysis, in TBS Buffer with 10% D_2_O. Data was processed using TopSpin 3.5 and Ccpnmr Analysis 2.4.2. ATPase and AK activity were checked by supplying ATP or ADP as substrate. For LptB2FG/C 5µM of complex was incubated with 5 mM of nucleotide and 1 mM MgCl_2_ in TBS buffer. When necessary, LptC_Δ[1-23]_ and LptA_Δ160_ were added at 10 μM. Batch experiments were incubated at 37°C for 7 h, flash frozen and transferred to 3 mm NMR tubes to be analyzed. 2 μM of LptB was incubated with 5 mM of nucleotide and 2.5 mM MgCl_2_ in TBS buffer. Batch experiments were incubated at 25°C for 17 h, flash frozen and transferred to 3 mm NMR tubes to be analyzed. STD experiments were recorded on a 700MHz cryoprobe at 25°C, with Bruker stddiffesgp.3 pulse sequence alternating on and off resonance at 0.5ppm and -40ppm, with a 60ms spinlock to suppress protein signals. 500µM of nucleotide (ADP or ADPβS) is mixed with 12.5 µM LptB (wt or mutant) with 50 µM MgCl2 in TBS buffer with 10 % Glycerol and 10 % D_2_O. Data are represented as Iref-Isat/Iref x 100. Error bars represent two times the standard deviation of the NMR experimental noise.

### Modeling of LptB_2_ with ADP

Modelling of ADP molecules to probe AK reaction site was done using the HADDOCK web server (35). LptB_2_ from *E. coli* in complex with ADP-magnesium was used [PDB code 4P32 (23)], and a second ADP docked, with an unambiguous restraint of 3.5±1Å between the 2 ADP β phosphates. A total of 1000, 200 and 200 structures were calculated rigid body, refinement and refinement in water respectively. Final structures produced two clusters of solutions.

## Acknowledgments

We thank A. Vallet and A. Favier for support with NMR. This work used the EM facilities at the Grenoble Instruct-ERIC Center (ISBG; UAR 3518 CNRS CEA-UGA-EMBL) with support from the French Infrastructure for Integrated Structural Biology (FRISBI; ANR-10-INSB-0005-02) and GRAL, a project of the University Grenoble Alpes graduate school CBH-EUR-GS (ANR-17-EURE-0003) within the Grenoble Partnership for Structural Biology. The IBS Electron Microscope facility is supported by the Auvergne Rhône-Alpes Region, the Fonds Feder, the Fondation pour la Recherche Médicale and GIS-IBiSA.

## Funding and additional information

This project was funded by the Train2Target project from the European Union’s Horizon 2020 Research and Innovation Program under the Marie Skłodowska-Curie grant agreement #721484.

## Supporting information

This article contains supplementary information.

