## supplementary information for "LptB_2_FG is an ABC transporter with Adenylate Kinase activity regulated by LptC/A recruitment"

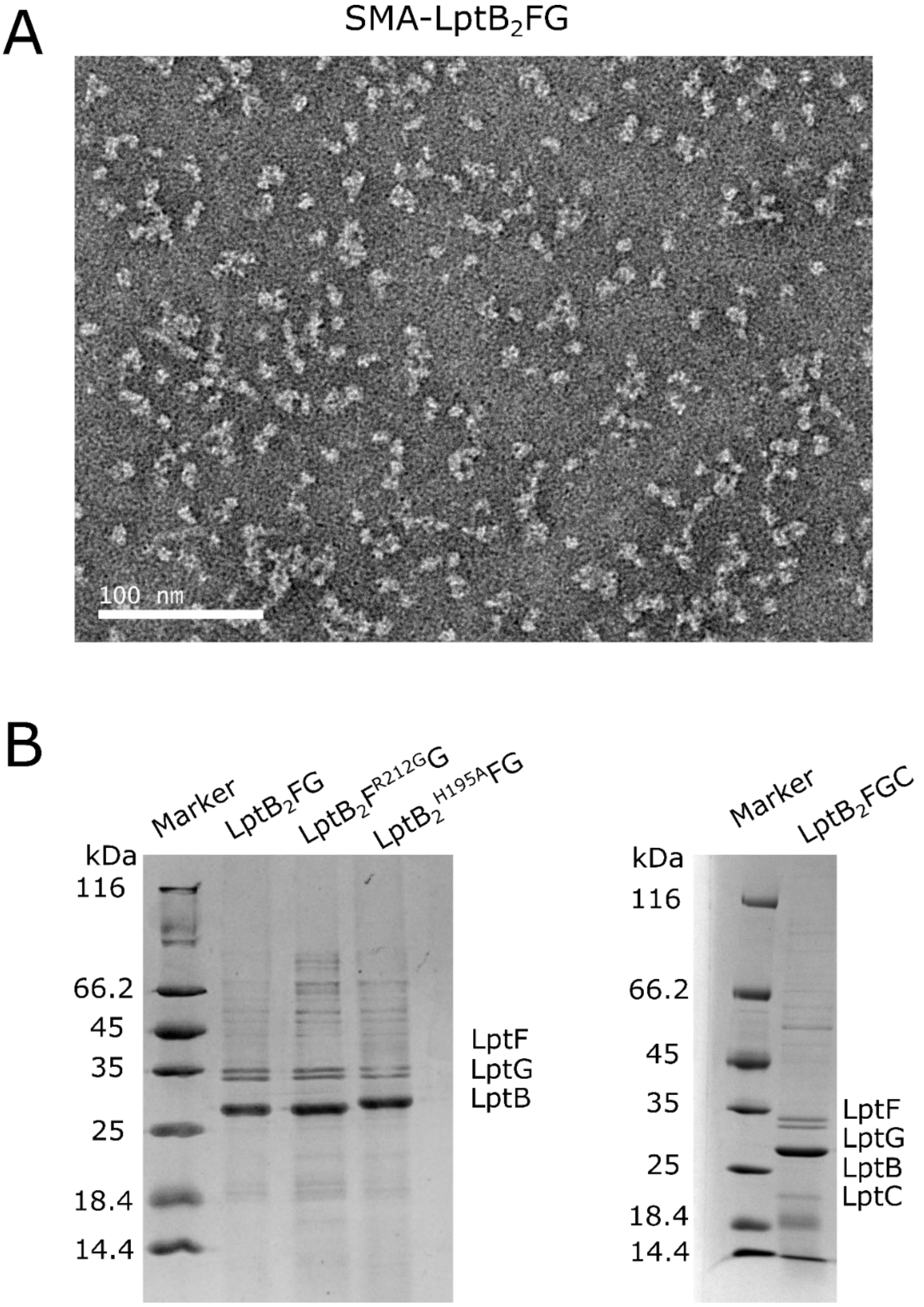


**Figure S1** – (A) Representative image of Electron Microscopy negative staining of SMA-LptB$_2$FG at 40µg/ml with Sodium Silico Tungstate staining; (B) SDS-PAGE 15% of LptB$_2$FG and LptB$_2$FGC complexes extracted into SMA nanodiscs. 2µg of complexes are injected.

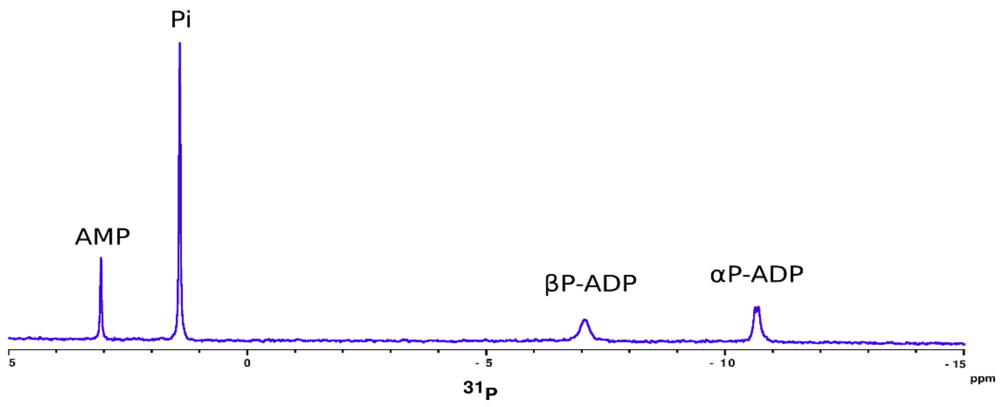


**Figure S2 –**1D $^{31}$P-NMR spectrum of LptB$_2$FG/ΔTM-C/A$_m$ after incubation with 5 mM ATP, with phosphorus resonances of ADP and AMP α-phosphate displayed.

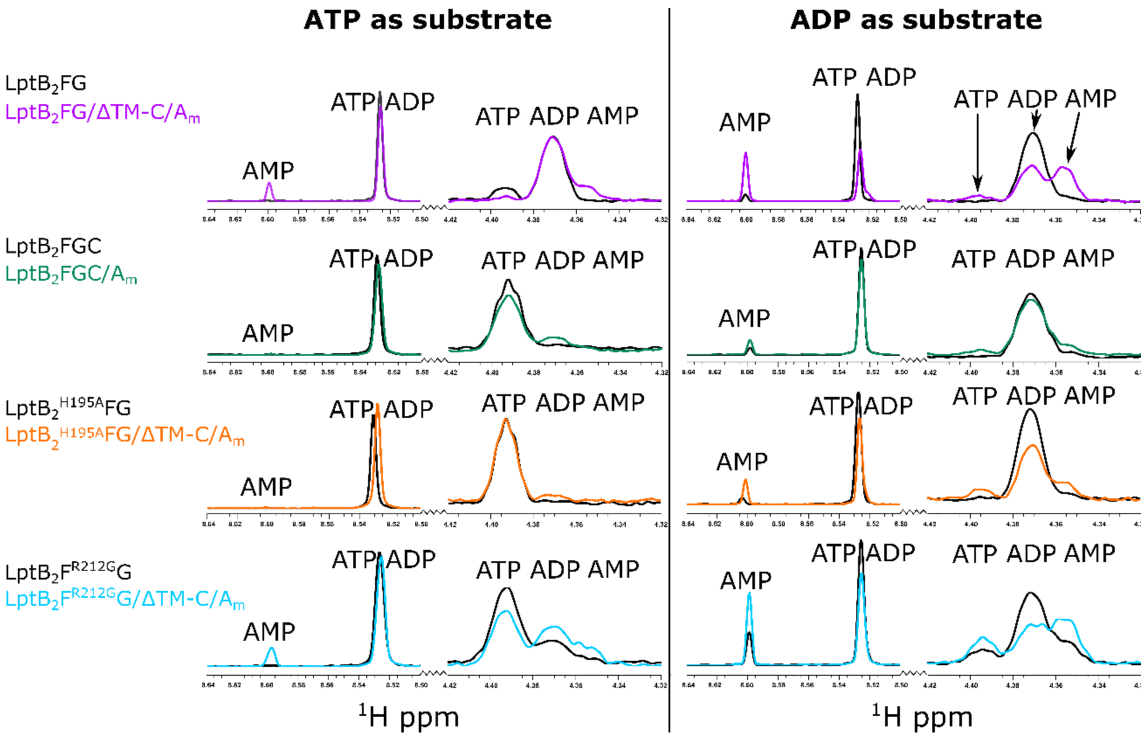


**Figure S3** – 1D $^{1}$H-NMR spectra showing ATPase/ AK activity of $LptB_2FG$, $LptB_2FG$/ΔTM-C/$A_m$, $LptB_2FGC/A_m$, $LptB_2^{H195A}FG$/ΔTM-C/Am and $LptB_2F^{R212G}G$/ΔTM-C/Am, either with ATP or ADP as substrate. Image displays zoom in the frequencies of the NMR probes used in the Adenosine H8 and Ribose H4' regions (spectrum level x4). Complex alone is displayed in black, while addition of remaining Lpt partners are displayed in colour-code accordingly, for both ATP-supplied and ADP-supplied experiments. H8 resonances from ADP and ATP are overlapped in these experimental conditions.

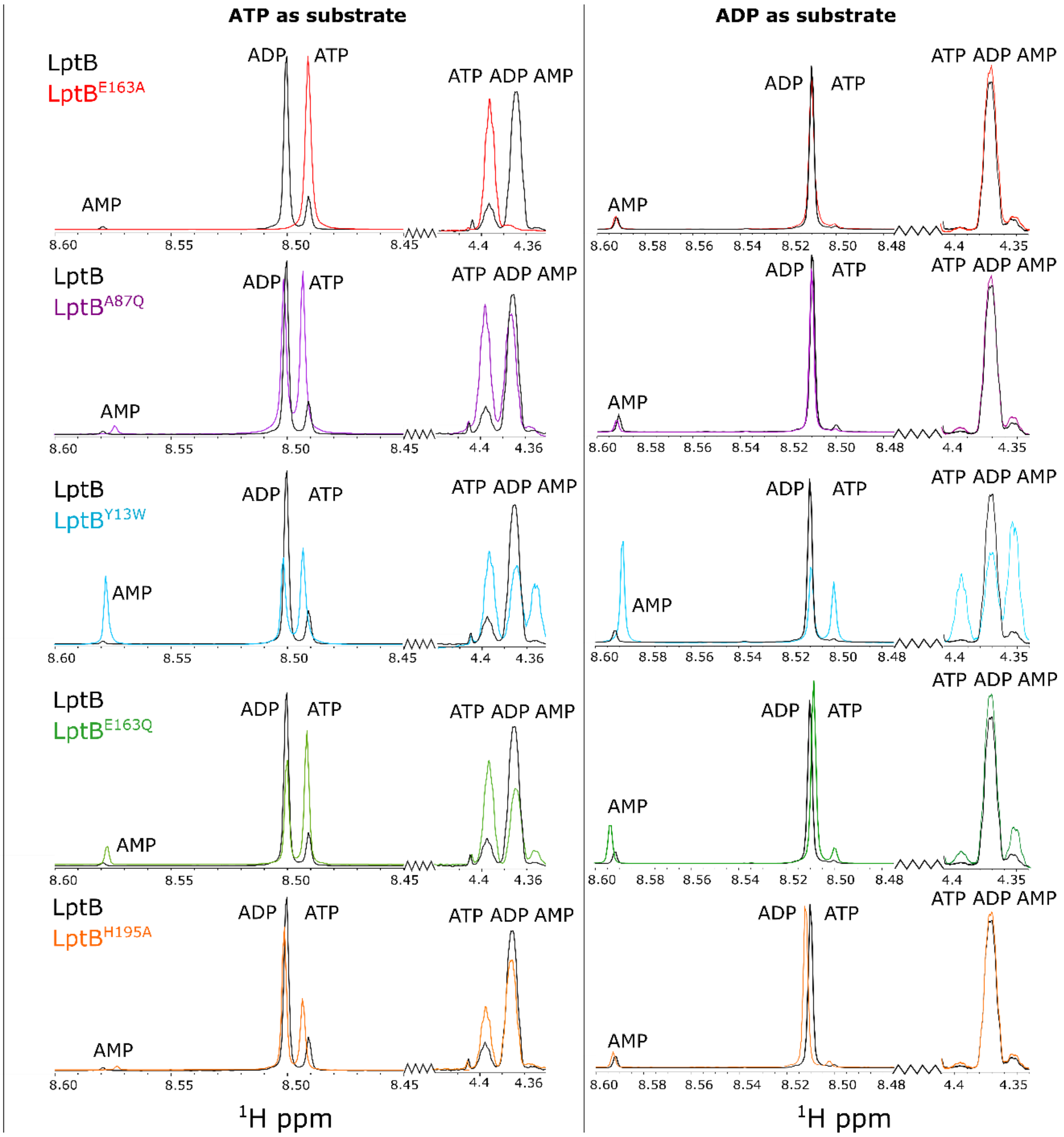


**Figure S4 –** 1D $^{1}$H-NMR spectra showing ATPase/ AK activity of isolated wild-type LptB and variant proteins, either with ATP or ADP as substrate. Image displays zoom in the frequencies of the NMR probes used in the Adenosine H8 and Ribose H4' resonances(spectrum level x4). Wild-type LptB protein spectrum is displayed in black, while each of the mutants are displayed in colour-code accordingly, for both ATP-supplied and ADP-supplied experiments.

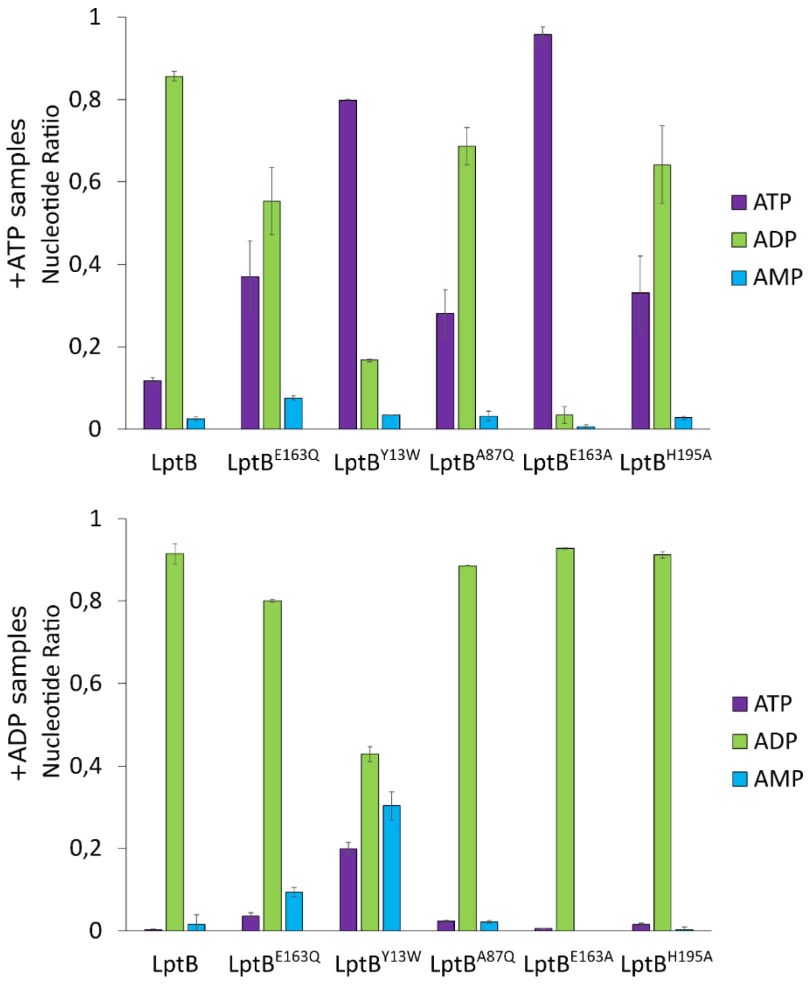


**Figure S5 –** ATPase and AK activities of $LptB_2$ wt, and variants in presence of either ATP or ADP, at 20°C. Nucleotide levels (ATP, ADP and AMP, in colour-code) were detected using a 1D $^{1}H$-NMR experiment in 3 mm tubes at 20°C, extracting peak intensities for each specie in 2 independent experiments.

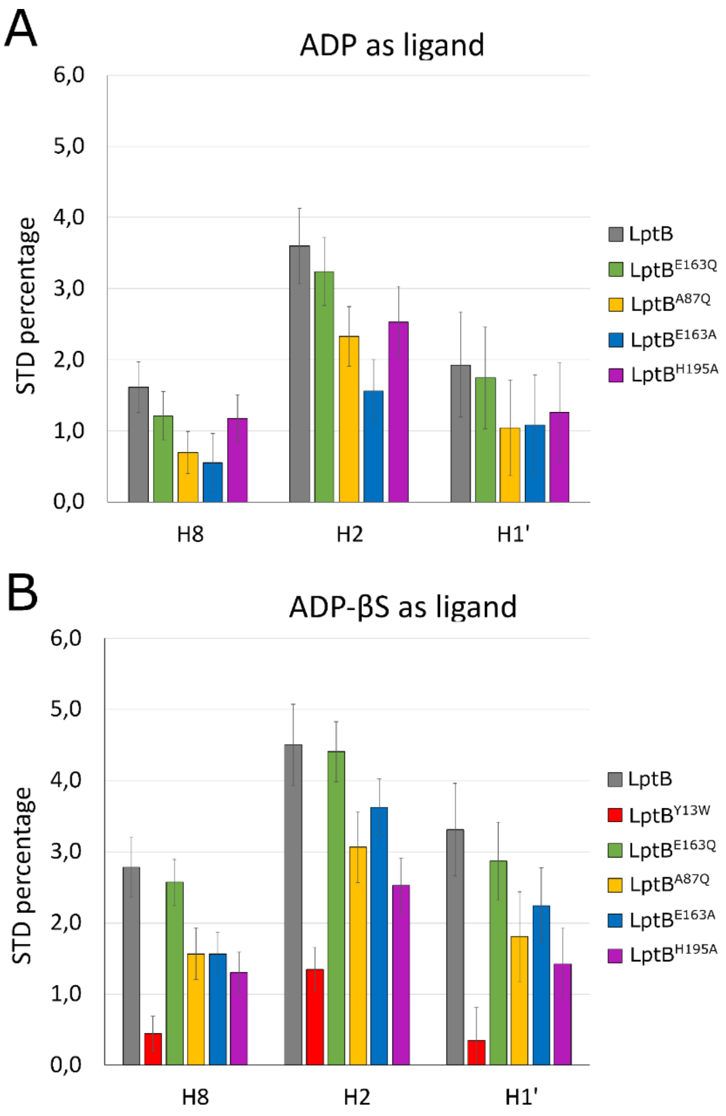


**Figure S6** – STD experiments using two ligands: (A) ADP and (B) ADP-βS, both for LptB proteins (wt and variants in colour-code). The three resonances shown are from H8, H2 and H1' on the adenosine of ADP and ADP-βS. Data are represented as (Iref-Isat/Iref). Error bars represent two times the standard deviation of the NMR experiments' noise.

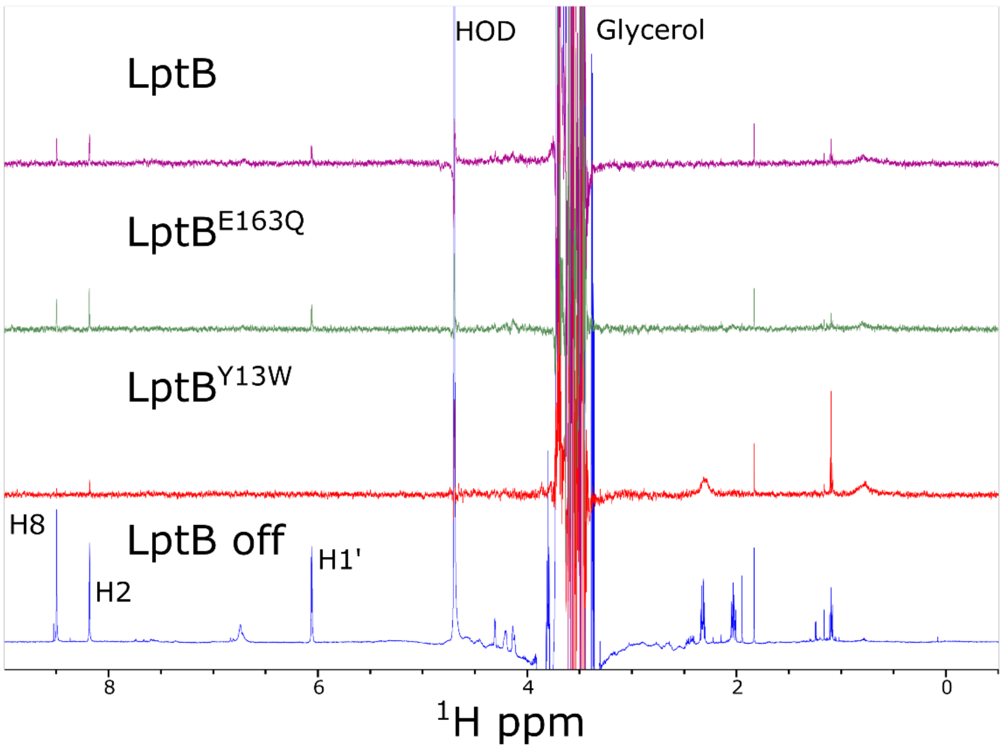


**Figure S7** -Saturation transfer difference spectra of LptB, LptB$^{E163Q}$ and LptB$^{Y13W}$ with ADPβS. (TOP) Difference spectra of ADPβS (OFF -ON resonance) in presence of LptB, LptB$^{E163Q}$ and LptB$^{Y13W}$.(Bottom) OFF resonance spectra of ADPβS in presence of LptB.

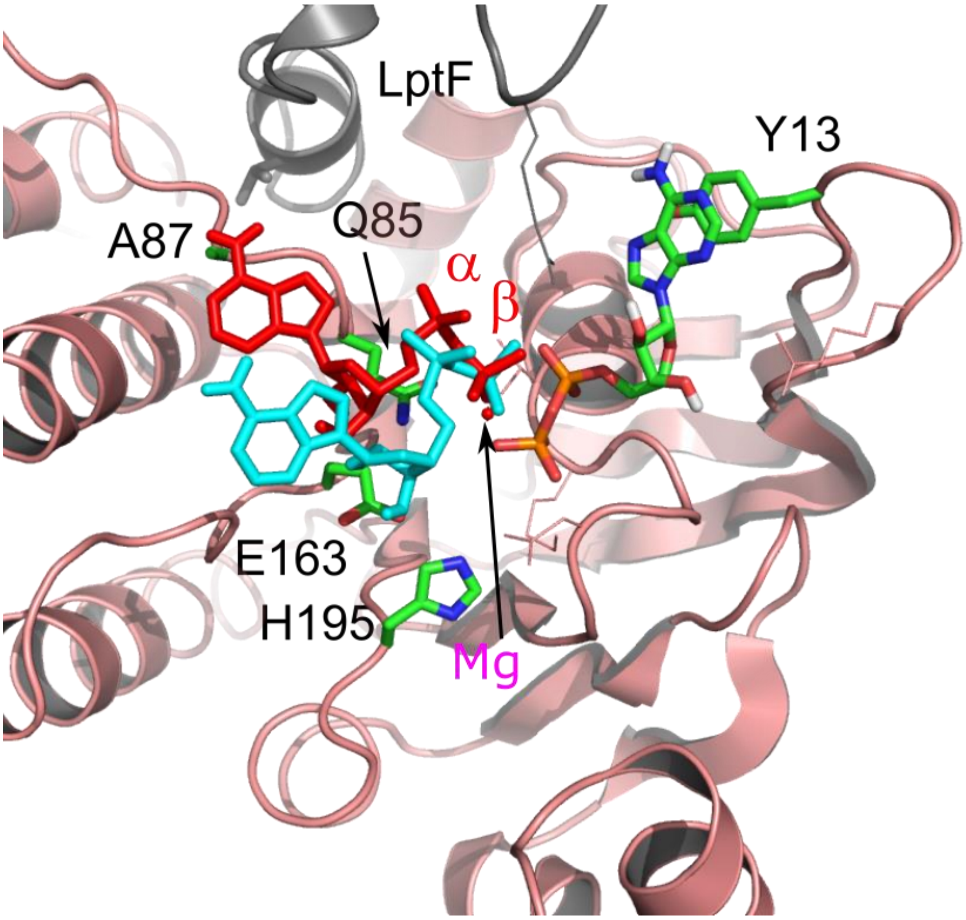


**Figure S8** – Haddock modelling of the LptB AK site. The best structures of the 2 main clusters of solutions were represented in red for the best cluster and in cyan for the second cluster on the 6S8G PSB structure, in the same orientation as Figure 3A. Magnesium is shown in magenta.

| | **Cluster 1** | **Cluster 2** |
|---|---|---|
| **HADDOCK score** | **-33.9 +/- 0.3** | **-26.4 +/- 1.2** |
| Cluster size | 173 | 27 |
| RMSD from the overall lowest-energy structure | 0.1 +/- 0.1 | 0.5 +/- 0.0 |
| Van der Waals energy | -13.0 +/- 0.4 | -10.9 +/- 0.7 |
| Electrostatic energy | -270.5 +/- 2.2 | -199.0 +/- 5.1 |
| Desolvation energy | 6.1 +/- 0.2 | 4.4 +/- 0.1 |
| Restraints violation energy | 0.1 +/- 0.04 | 0.1 +/- 0.01 |
| Buried Surface Area | 444.8 +/- 3.5 | 355.0 +/- 3.4 |
| Z-Score | -1.0 | 1.0 |

Table S1: Haddock statistics of the two cluster of solutions obtained in the docking procedure.
